# Evidence for unfamiliar kin recognition in vampire bats

**DOI:** 10.1101/2019.12.16.874057

**Authors:** Simon P. Ripperger, Rachel A. Page, Frieder Mayer, Gerald G. Carter

**Author notes:** Correspondence: Simon Ripperger, Gerald Carter.

## Abstract

Kin discrimination allows organisms to preferentially cooperate with kin, reduce kin competition, and avoid inbreeding. In vertebrates, kin discrimination often occurs through prior association. There is less evidence for recognition of unfamiliar kin. Here, we present the first evidence of unfamiliar kin recognition in bats. We captured female vampire bats (*Desmodus rotundus*) from a single roost, allowed them to breed in captivity for 22 months, then released 17 wild-caught females and six captive-born daughters back into the same wild roost. We then used custom-built proximity sensors to track the free-ranging social encounters among the previously captive bats and 27 tagged control bats from the same roost. Using microsatellite-based relatedness estimates, we found that previously captive bats preferentially associated with related control bats, and that captive-born bats preferentially associated with unfamiliar kin among control bats. Closer analyses showed that these unfamiliar-kin-biased associations were not caused by mothers or other familiar close kin, because the kinship bias was evident even when those bats were not nearby. This striking evidence for unfamiliar kin recognition in vampire bats warrants further investigation and provides new hypotheses for how cooperative relationships might be driven synergistically by both social experience and phenotypic similarity.

## Background

Genetic relatedness plays a pivotal role in the evolution of social behaviour [1, 2]. For organisms that regularly interact with relatives and non-relatives, kin discrimination allows for social behaviours that increase inclusive fitness, including helping kin, reducing kin competition, and avoiding inbreeding. In vertebrates, kin discrimination often occurs through prior association [3], but there are far fewer reports for the ability to detect unfamiliar kin, which requires some form of phenotype matching [4]. Experimental evidence for unfamiliar kin recognition comes from fish [5], amphibians [6, 7], birds [8], rodents [9, 10], and primates [11, 12].

Surprisingly little is known about kin discrimination in bats. Many of the more than 1,400 bat species demonstrate traits linked to the evolution of kin discrimination, including a low mean and high variance in group kinship [13, 14], budding dispersal [15, 16], preferred associations [14], and in some cases, nepotistic helping that bestows large fitness benefits [17]. The clearest example of kin-biased helping in bats is regurgitated food sharing in vampire bats [17]. Common vampire bats show preferred co-roosting associations with both kin and nonkin (mean within-roost relatedness = 0.08 [18, 19]). Yearling males disperse whereas females are typically philopatric and form long-term cooperative relationships within and between matrilines [17, 20]. Food-sharing in the wild is kin-biased even when controlling for co-roosting association [17]. However, in all studies to date with this species, cooperating kin have also been familiar, so it remains unclear whether vampire bats can identify unfamiliar kin.

Here, we show that vampire bats (*Desmodus rotunds*) preferentially associated with kin that they have been separated from for almost two years, and using captive-born bats, we also show evidence for unfamiliar kin recognition. We captured female vampire bats from a wild population, housed them in captivity for 22 months, then attached proximity sensors on 17 adult females and six captive-born daughters, and released these 23 ‘test bats’ back into the roost where they—or their mothers—were originally captured. As a control group, we also fitted a sample of 27 ‘control bats’ from the same wild population with the same proximity sensors, which track the time, duration, and signal strength (as a distance estimate) of all tagged bats within proximity. By genotyping the tagged bats, we were able to test for kin-biased association between the test bats and the control bats. Surprisingly, the captive-born bats spent more time near unfamiliar kin in the control group than expected by chance, even when accounting for the presence of familiar kin. This finding is the first evidence of unfamiliar kin recognition in a bat.

## Methods

### Proximity sensing

To track free-ranging associations between the 23 previously captive test bats and 27 control bats, we fitted them with 1.8-g proximity sensors, custom-developed for the BATS tracking system [21, 22]. The animal-borne sensors sense dyadic proximity among all tagged individuals that are within communication range (up to five meters), every two seconds. When two sensors come within communication range, the start of a meeting is logged. When the sensors go outside this range for 10 or more seconds, the meeting is closed and the on-board memory stores the meeting start time, duration, maximum received signal strength indicator (RSSI), and both sensor IDs. All meeting data are remotely downloaded to a base station, which we placed inside the bat roost during this study [23]. The tagged bats weighed 27-48 g, so tags were 3.8-6.7 % of body mass.

### Study design

To create the test group, we captured adult females using mist nets outside a hollow tree in Tolé, Panama on December 13, 2015, then housed them in a captive colony in a 1.7 × 2.1 × 2.3 m outdoor flight cage at the Smithsonian Tropical Research Institute in Gamboa, Panama, for 22 months [23, 24]. During that time, six female offspring were born into the captive colony, which were 10-19 months old at the time of our study. The bats in this ‘test group’ were individually marked with subcutaneous passive integrated transponders and a visually unique combination of forearm bands.

On September 20, 2017, we fitted the 23 test bats with proximity sensors and released them at the original hollow tree at 2012 h. As a control group, we also captured, banded and fitted proximity sensors to 27 adult females caught at the same roost and released them back between 0450 h to 0630 h on September 20. The hollow tree was large enough to contain distinct social groups, and we estimated the tree to contain about 200 vampire bats based on captures and photographs. The main cavity was about 1.5 m wide and 2.5 m high, and several smaller cavities branched off from the main one.

To define roosting association rates at a given time period, we summed durations of meetings with maximum RSSI that correspond to at least 50 cm of proximity, following a previous analysis [23]. We measured dyadic associations from 0600 h to 2400 h from September 20-28.We did not include the hours between 0000 h and 0600 h because many individuals went foraging during this time period. Two control bats and one wild-born test bat left the study site on the first sampling day. Relatedness data was missing for one control bat, and one captive-born bat was unrelated to all others in the wild colony. Consequently, the data underlying the analyses come from 21 test bats (including five captive-born daughters) and 24 control bats.

### Genetic relatedness

To measure relatedness, we extracted DNA from a 3-4 mm wing biopsy punch in 80% or 95% ethanol using a salt–chloroform procedure, then used a LI–COR Biosciences DNA Analyzer 4300 and the SAGA GT allele scoring software to genotype individuals at 17 polymorphic microsatellite loci (see Table S2 in [23]). Allele frequencies were based on 100 bats from Tolé and nine bats from another site, Las Pavas, Panama. Genotypes were 99.9% complete. We used the Wang estimator in the R package ‘related’ [25] to obtain an initial kinship estimate based on relatedness, then we assigned a zero kinship to dyads with negative relatedness estimates to ensure that any kin-biased associations were not driven by negative relatedness values. We also assigned a kinship of 0.5 for known mother-offspring dyads or dyads with relatedness estimates greater than 0.5. We detected no significant differences in the mean pairwise relatedness between two control bats (mean = 0.077 [95% CI = 0.064 - 0.088], n = 276), and a test and control bat (mean = 0.067 [95% CI = 0.059 - 0.076], n = 552), as would be expected from a random sample of bats captured from the same colony.

### Data analysis

We first tested for kin-biased associations across all bats and days (see supplement for methods and results). Next, we tested for kin-biased associations only in test-control bat dyads. To do this, we calculated the Pearson’s correlation between relatedness and association rates in test-control dyads separately for each day, and then calculated the mean daily correlation as the effect size. We compared this effect size to the expected null distribution of effect sizes generated from network permutations of the bats present in the tree within each day, following [23]. This permutation procedure controls for the presence or absence of bats on each day. To estimate a mean effect size and 95% confidence interval (CI) for each test bat, we bootstrapped (5,000 iterations) the effect size across the possible test-control dyads within each test bat. This allowed us to visualize each bat’s contribution to the overall kin-biased association.

To test for evidence of unfamiliar kin recognition, we analysed association rates between captive-born test bats and control bats that were previously unfamiliar. One challenge here is that captive-born bats know their mothers (and possibly other familiar kin), and these familiar kin could in effect ‘introduce’ or bias the association between a captive-born bat and an unfamiliar relative in the wild colony. For example, if captive-born bats stay close to their mothers, and those mothers associate more with related control bats (also related to her offspring), the result would be the *false* appearance of kin-biased association between the captive-born test bats and unfamiliar control bats. Therefore, to remove this possibility, we first looked at whether the unfamiliar kin encounters were initiated by the mothers of captive-born bats, by plotting the temporal sequence of encounters (hourly nonzero association rates) for captive-born daughters and their mothers with each control bat. Next, we conducted a permutation test which removed the effect of mothers. We calculated hourly association times between a captive-born bat and all possible control bats, and then removed all hourly association times where the captive-born bat’s mother was associated with the control bat at any time during that same hour. These ‘filtered’ associations represent the time each captive-born bat spent with each control bat without the mother nearby. For an effect size, we summed these hourly association times within each unfamiliar dyad and calculated their correlation with dyadic kinship. To get a distribution of expected effect sizes under the null hypothesis, we repeated this same procedure after first randomizing each captive-born bat’s association times across control bats within each hour, while keeping the mother’s associations the same (5,000 randomizations).

We then repeated this analysis, but instead of only removing the hours when mothers were nearby, we removed hours when any familiar close kin of the captive-born bat was nearby (with close kin defined as a relatedness estimate of 0.125 or higher). Both permutation tests detected the same effect when we conducted an even more conservative test that only included non-zero association rates in the analysis (i.e. testing the effect of kinship on encounter duration but not probability).

## Results

We detected a higher-than-chance correlation between genetic relatedness and association rates (i.e. kin-biased association) across all bats (see supplement). When focusing only on test-control dyads we also found that association rates between formerly captive test bats and control bats correlated stronger with genetic relatedness than expected by chance (r = 0.091, p < 0.0002; Figure 1a), and this was true even when excluding captive-born bats (r = 0.076, p < 0.0002). To test for kin-biased association between unfamiliar bats, we limited our analysis to associations in unfamiliar dyads of a captive-born bat and a control bat in the absence of familiar kin. Captive-born bats associated with unfamiliar kin more than expected by chance, even when controlling for the presence of the captive-born bats’ mother (r = 0.19, p = 0.002; Figure 1b) or the presence of other familiar close kin (r = 0.15, p = 0.0004; Figure 1c).

**Figure 1.**
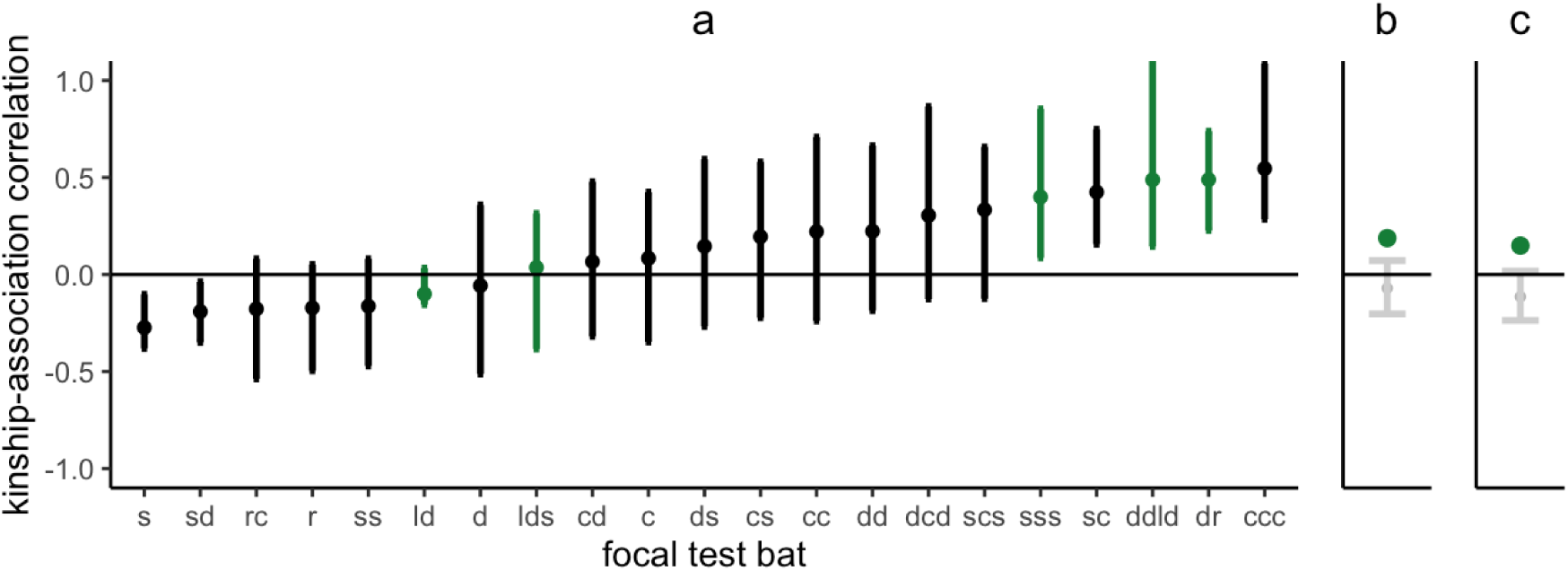
Kin-biased associations. Mean correlations between kinship and association with control bats, shown for each of the 21 test bats (a), were biased above zero. Error bars are 95% CIs from bootstrapping across the 24 control bats. Kin-biased association in captive-born bats (green) were evident even after removing encounters that involved the captive-born bat’s mother (b) or any familiar kin (c). Larger green dots shown mean correlations across captive born bats and grey error bats show the 5% and 95% quantiles for the expected mean correlation to reflect a one-sided test. The timing of kin associations are shown in supplementary Figures S1–S5. The slopes and values of pairwise kinship and association for each bat are shown in supplementary Figure S6.

## Discussion

High-resolution proximity sensing revealed the first evidence of unfamiliar kin recognition in a bat. When we released captive female vampire bats back into the wild, they preferentially roosted near closer kin that they were separated from for almost two years. More importantly, five captive-born daughters with genetic relatives in the control group preferentially associated with unfamiliar kin, even when the mothers or other familiar close kin were not present nearby. The chronology of social encounters (Figure S1–5) suggests that mothers did not initiate most of these encounters. For example, the mother was not nearby in eight of the eleven sampled cases where the captive-born daughter encountered unfamiliar kin from the control group (relatedness 0.125 or higher) for the first time.

Vampire bat social bonds are driven by both kinship and past social experience [17, 23, 24, 26–29]. Both food sharing and association is kin-biased [17], but even when controlling for kinship, food sharing is reciprocal [24, 26–29] and the previously captive bats in this study also preferentially associated in the wild with the bats that more frequently groomed or fed them in captivity [23]. Furthermore, playback studies show that captive female vampire bats are more attracted to the contact calls of unrelated food donors than to the contact calls of related groupmates that were non-donors [26]. In this study, we show evidence that kinship can also drive association without past social experience. The ability to recognize unfamiliar kin could allow for female bats to preferentially associate or cooperate with paternal kin or avoid inbreeding with related males.

Kinship alone, however, is clearly not sufficient for social integration. Although the captive-born bats associated more with relatives, they appeared to fail to integrate into the wild roost [23]. All the captive-born bats left the roost by day six and we did not see them return. Bites marks on some captive-born bats suggest aggression played a role in their departure (see [23] for details).

Recognizing unfamiliar kin suggests some form of phenotype matching, where kinship cues (e.g. olfactory [30], visual [11, 12], acoustic [31], or multimodal [32]) are either learned from familiar kin or matched to one’s own phenotype [5]. In vampire bats, both acoustic and olfaction cues are plausible candidates for unfamiliar kin recognition. Common vampire bats possess an intact vomeronasal system for detecting pheromones, and have at least twice as many intact vomeronasal type-1 receptor genes as other sampled bats [33, 34]. Vocal phenotype matching is also plausible given evidence in primates (e.g. [31]) and the primacy of sound in the social lives of bats [35]. Future work in this species should test whether calls of unfamiliar kin can be discriminated in playback experiments and whether unfamiliar kin are more likely to develop food-sharing relationships.

Our findings relied on extracting signatures of novel behaviours from a large high-resolution dataset enabled by recent and revolutionary advances in biologging [21], in this case, miniaturized proximity sensors that log social encounters within largely inaccessible sites [23, 36]. We suspect that ongoing advances in biologging technology will continue to reveal many new insights into species that are difficult to directly observe [37–39].

### Ethics

All experiments were approved by the Smithsonian Tropical Research Institute Animal Care and Use Committee (#2015-0915-2018-A9 and #2017-0102-2020) and by the Panamanian Ministry of the Environment (#SE/A-76-16 and #SE/AH-2-17).

## Acknowledgements

We thank O. Castrellón for permission to conduct field work on his property, and V. Flores, M. Le Chevallier, M. Nowak, D. Josic, J. Berrío-Martínez, G. Cohen, B. Cassens, and N. Duda for their assistance. We are grateful to I. Waurick for her valuable assistance and expertise during molecular lab work.

## Supplementary Information

### Kin-biased association across all bats

We tested for kin bias in association rates among all test and control bats summed over the entire study period (including test-test, control-control, and test-control dyads). To do this, we used the multiple quadratic assignment procedure with double semi-partialling in the asnipe R package [S1]. This network permutation test allows us to test the effect of kinship while controlling for a covariate [S2]. Given that test bats preferred to associate with each other [S3], we controlled for this assortativity by including test-test dyad type as a covariate (test-test dyads = 1, other dyads = 0) in the first permutation test and including test-control dyad type (test-control dyads =1, other dyads = 0) in the second permutation test. Across all bats and days, we detected that more closely related bats were associated longer than expected by chance (controlling for test-test dyad type: kinship β = 0.176, p < 0.0002; controlling for test-control dyad type: kinship β = 0.163, p < 0.0002).

### Figures S1–S5: Chronologies of encounters among unfamiliar kin (i.e. captive-born test bats and wild control bats)

**Figure S1:**
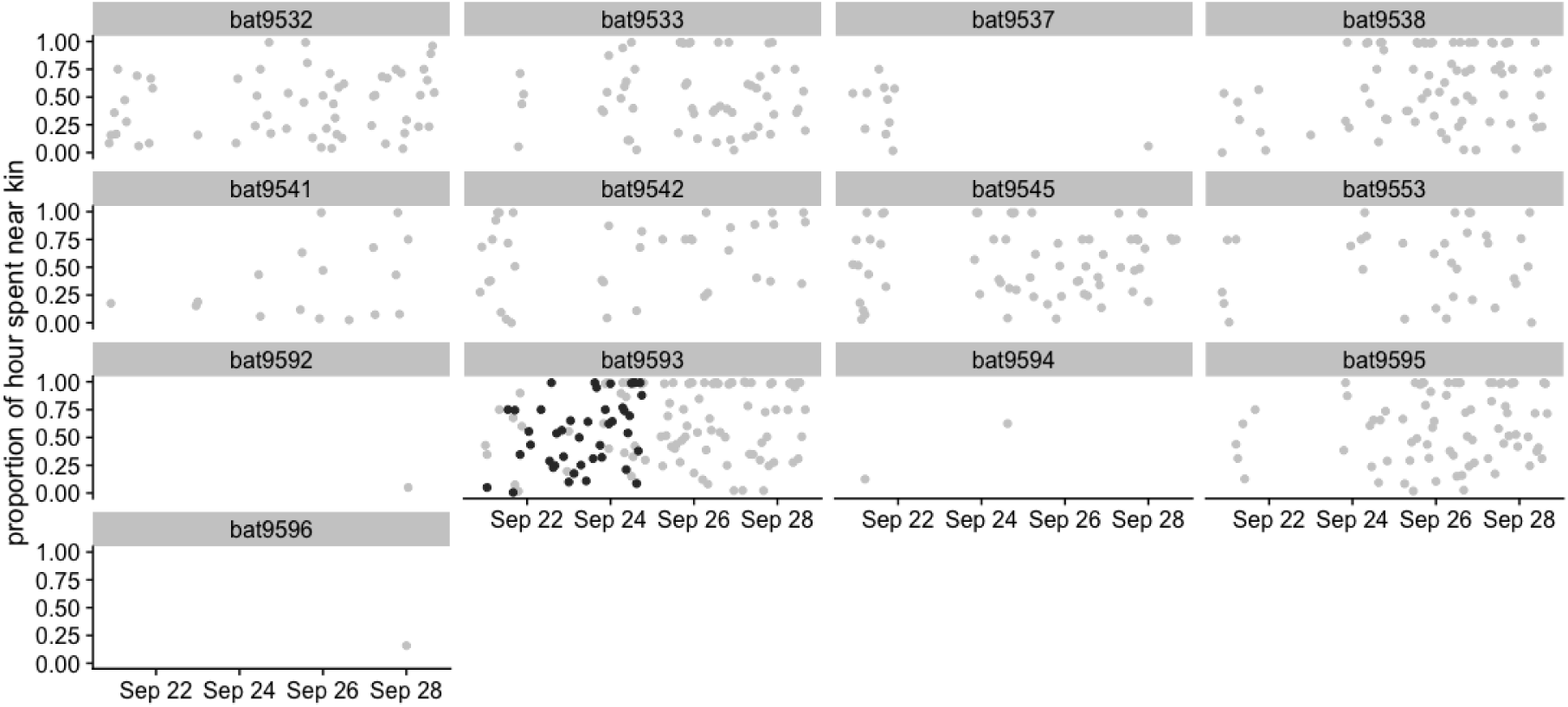
Hourly association rates with wild kin (all control bats with genetic relatedness greater than 0.124 to bat SSS) are shown for bat SSS (captive-born; black dots) and for its mother (wild-born formerly-captive bat; grey dots).

**Figure S2:**
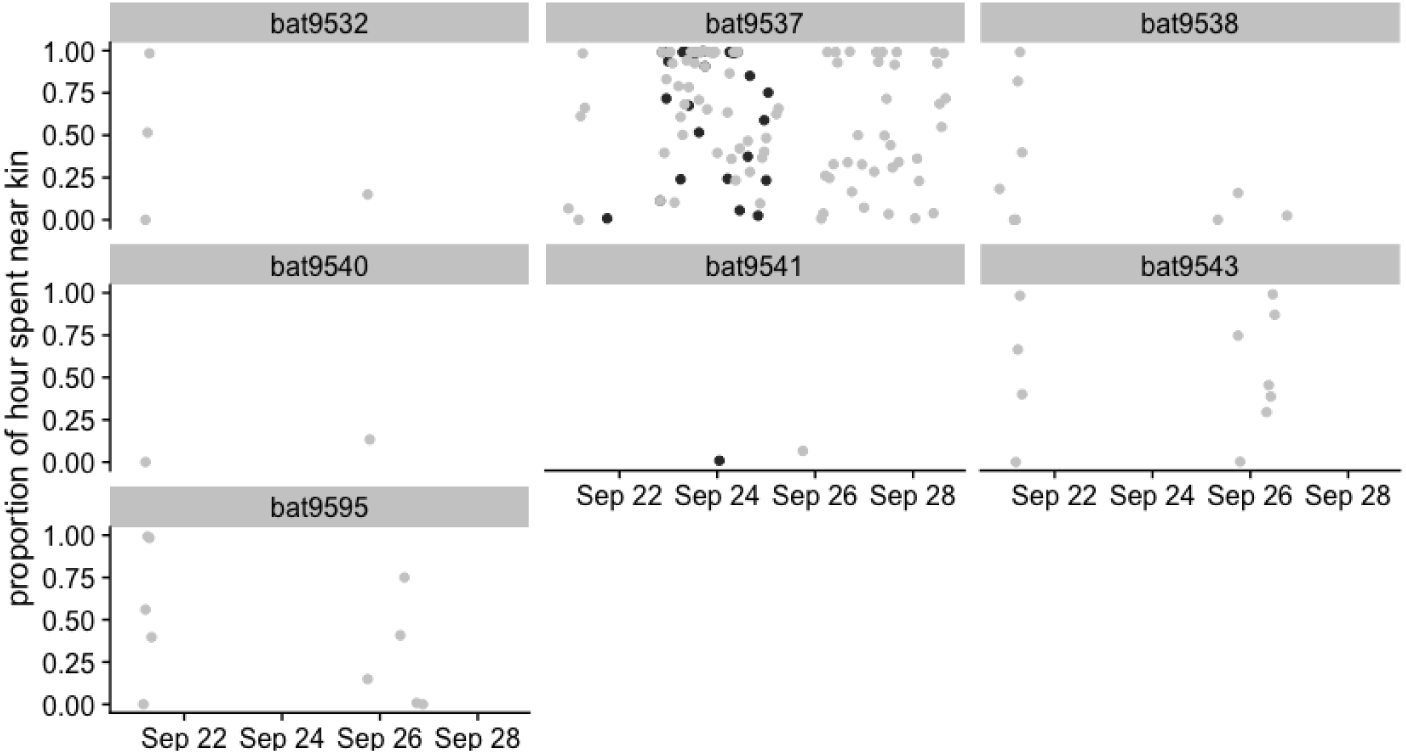
Hourly association rates with wild kin (all control bats with genetic relatedness greater than 0.124 to bat DDLD) are shown for bat DDLD (captive-born; black dots) and for its mother (wild-born formerly-captive bat; grey dots).

**Figure S3:**
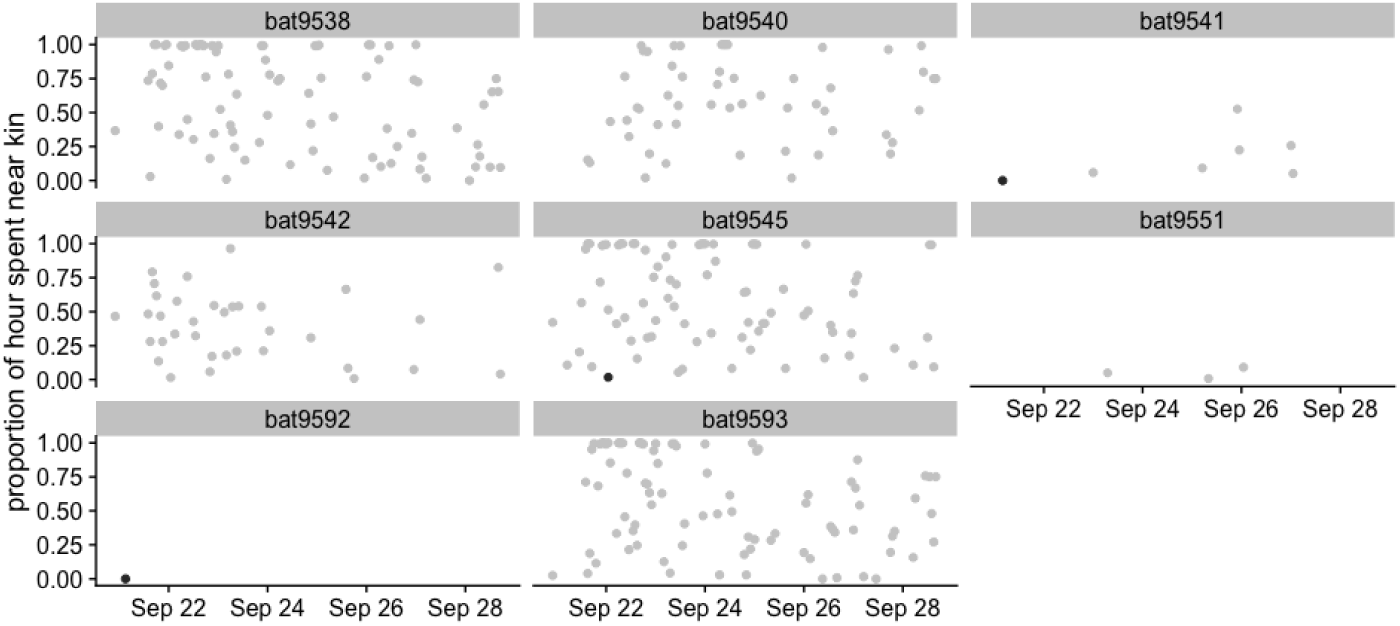
Hourly association rates with wild kin (all control bats with genetic relatedness greater than 0.124 to bat LD) are shown for bat LD (captive-born; black dots) and for its mother (wild-born formerly-captive bat; grey dots).

**Figure S4:**
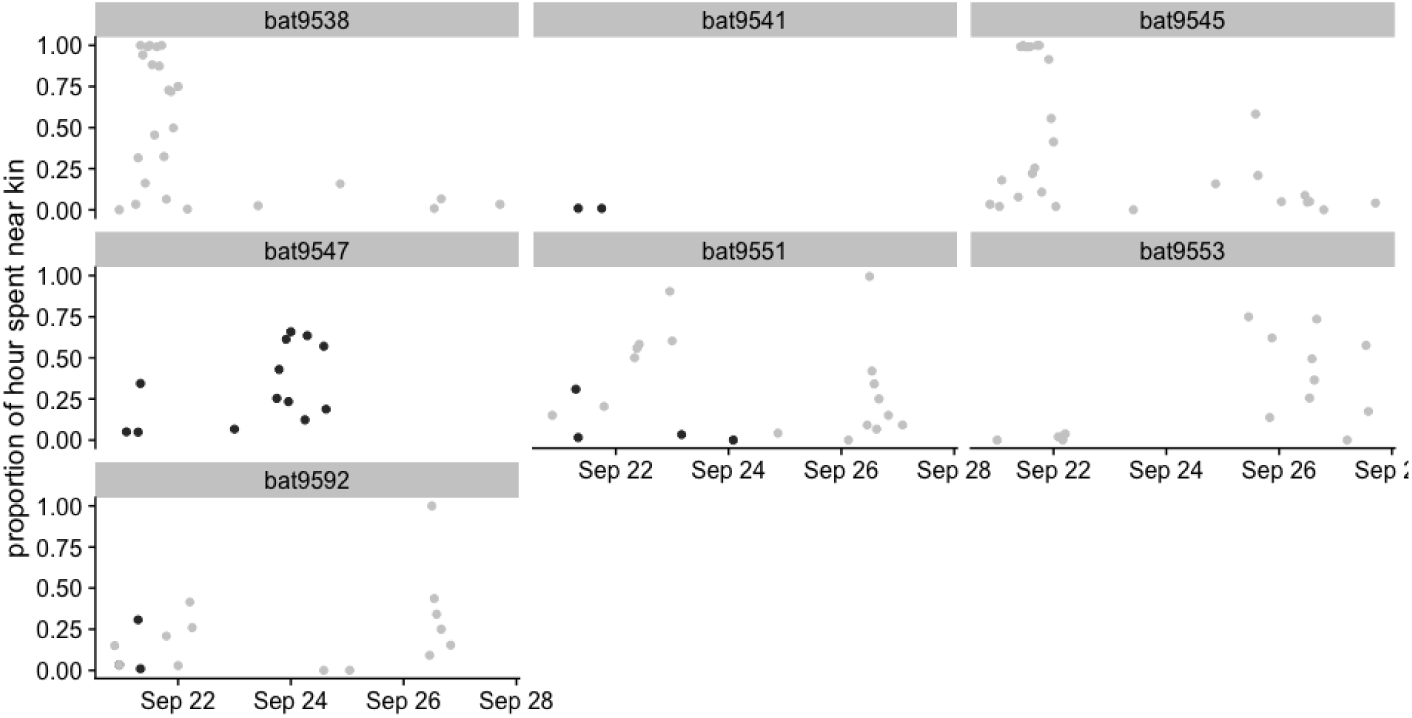
Hourly association rates with wild kin (all control bats with genetic relatedness greater than 0.124 to bat LDS) are shown for bat LDS (captive-born; black dots) and for its mother (wild-born formerly-captive bat; grey dots).

**Figure S5:**
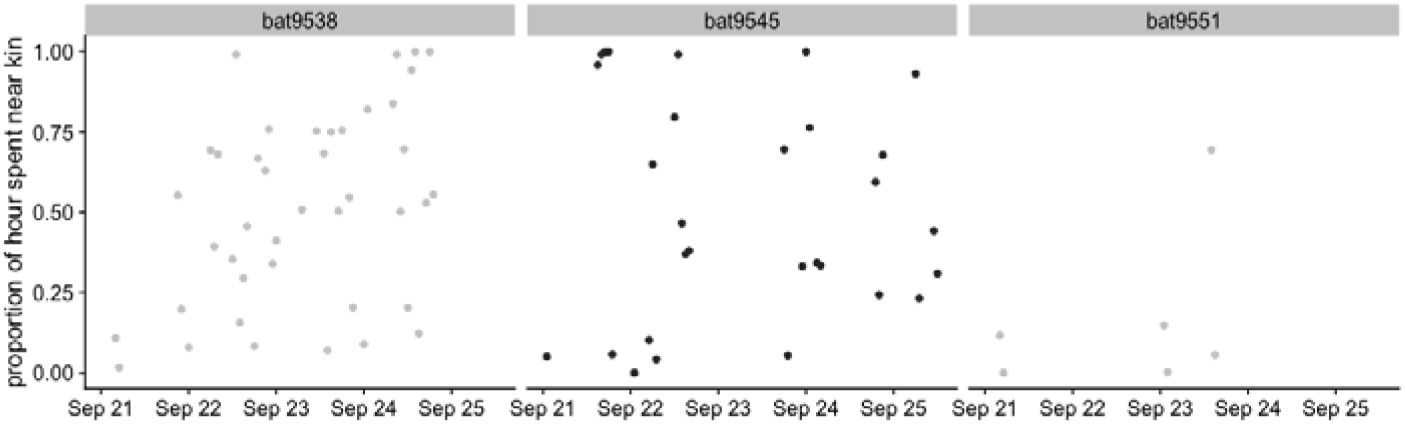
Hourly association rates with wild kin (all control bats with genetic relatedness greater than 0.124 to bat DR) are shown for bat DR (captive-born; black dots) and for its mother (wild-born formerly-captive bat; grey dots).

**S6.**
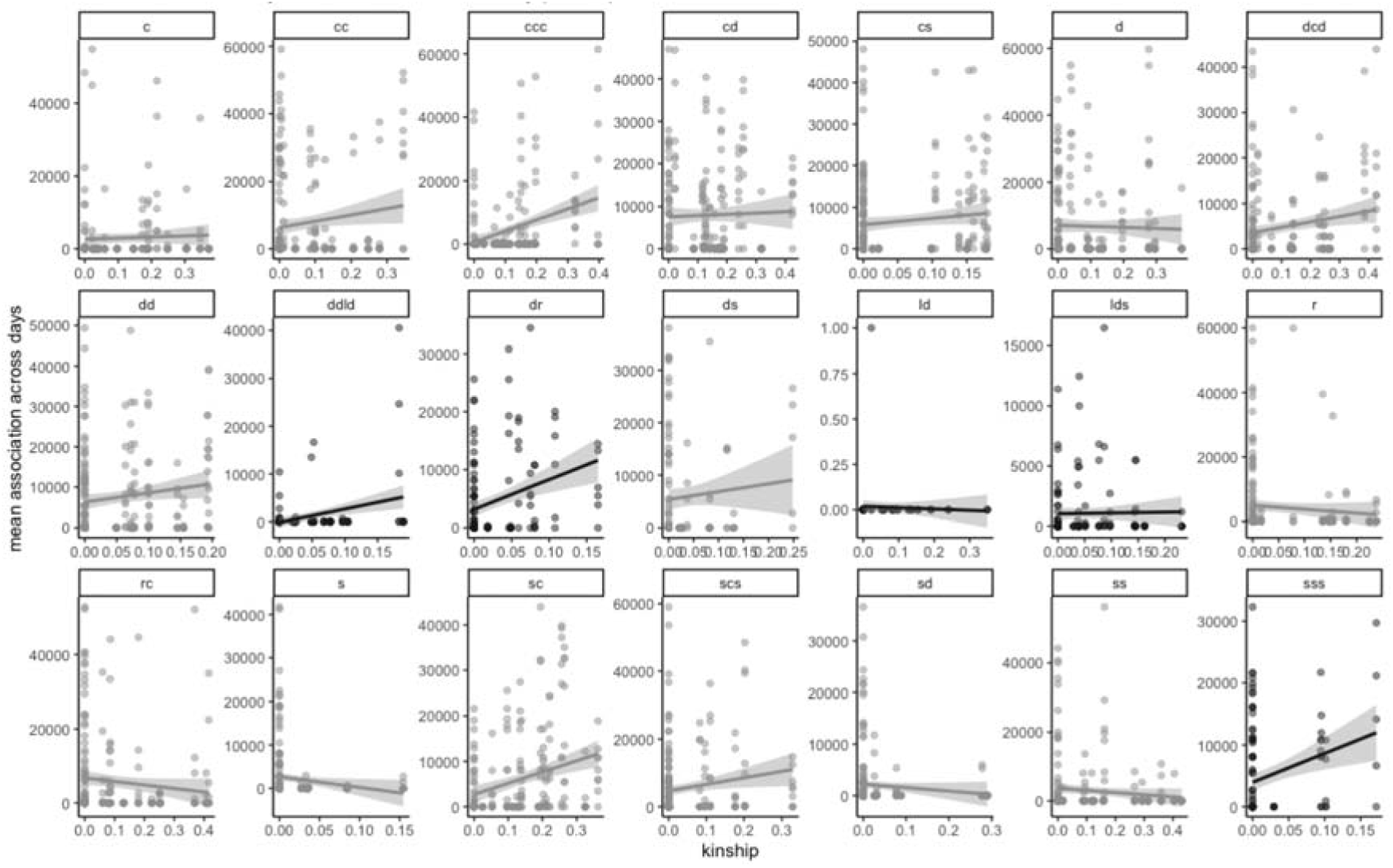
Kin-biased associations by bat. Each panel is a test bat. Each data point (n = 3199) represents an association rate (s per sample day) between that test bat and a control bat on a given sampling day. Points are transparent to show overlapping points. Black slopes are captive-born test bats and grey slopes are wild-born test bats. Slopes and their 95% confidence intervals are estimates by treating association rates on different days as independent (because bats left the tree and returned between sample days).

